# A pilot investigation of chemonucleolysis-induced intervertebral disc degeneration in the ovine lumbar spine

**DOI:** 10.1101/384743

**Authors:** Ryan Borem, Joshua Walters, Allison Madeline, Lee Madeline, Jeremiah Easley, Sanjitpal Gill, Jeremy Mercuri

**Author notes:** Corresponding Author*: (JM).

## Abstract

Intervertebral disc (IVD) degeneration (IVDD) initiates in the nucleus pulposus (NP) and is marked by elevated levels of pro-inflammatory cytokines and matrix-degrading proteases, leading to structural and functional disruption. IVDD therapeutics are currently being investigated; however, such approaches require validation using large animal models that recapitulate clinical, biochemical, and biomechanical hallmarks of the human pathology. Others have previously utilized intradiscal administration of chondroitinase-ABC (C-ABC) to initiate IVDD in the NP of sheep lumbar IVDs. While these studies examined changes in IVD height, hydration, and tissue micro-architecture, changes in biochemical content and mechanical properties were not assessed. Thus, the objective herein was to comprehensively characterize this ovine model IVDD for salient features reported in human degenerate IVDs by evaluating biochemical, biomechanical, and histological changes. Briefly, C-ABC (1U) was administered via intradiscal injection into the L_1/2_, L_2/3_, and L_3/4_ IVDs, and degeneration was assessed at 6- and 10-weeks via longitudinal magnetic resonance (MR) imaging. After 6 weeks, degenerative samples showed significant reductions in IVD heights (p=0.048) and MR imaging index (p=0.048), which worsened at 10 weeks. Post-mortem degenerate and controls IVDs were evaluated for differences in interleukin-1β concentration, axial and torsional functional spinal unit kinematics, and histological microarchitecture. Degenerate IVDs demonstrated significantly elevated concentrations of interleukin-1β (p=0.002). Additionally, degenerative samples showed increased creep displacement (p=0.022) and compressive stiffness’s (p=0.007) concurrent with decreased long-term elastic (p=0.007) and viscous dampening coefficients (p=0.002). Histological analysis of degenerative IVDs showed changes in microarchitecture, including derangement of the nucleus pulposus and annulus fibrosus tissue as well as cartilaginous end-plate irregularities. This pilot study demonstrated that intradiscal injection of 1U C-ABC induces significant and progressive degeneration of sheep lumbar IVDs over the time course investigated. The changes observed in this pilot’s study small sample size resemble the hallmarks of moderate to severe IVD degeneration observed in humans. Further study is warranted on a larger sample size to further validate these findings.

## Introduction

The intervertebral disc (IVD) is a fibro-cartilaginous structure adjoining vertebral bodies of the spine. The primary role of the IVD is to support and transmit axial loads, while allowing for spinal mobility and stability during activities of daily living. Each IVD comprises three morphologically distinct regions. The centralized core of the IVD is known as the nucleus pulposus (NP); a hydrophilic, matrix composed of aggrecan and collagen type II. The NP is circumferentially sequestered by the annulus fibrosus (AF); a ring-like structure consisting of 15-25 concentric layers of type I collagen whose fiber preferred direction is oriented at ± 28-43° to the transverse axis of the spine in alternating layers.[1] This results in a fiber-reinforced composite structure with an ‘angle-ply’ microarchitecture. The third region of the IVD is known as the cartilaginous end-plate (CEP) which is a thin layer of hyaline cartilage that is located between the inferior and superior surfaces of the IVD and adjacent vertebral bodies. The CEP serves as a mechanical barrier and allows for nutrient transport between the IVD and adjacent vertebral bodies.[2]

Approximately 1.5 to 4 million adults in the U.S. have IVD-related low back pain (LBP), leading to lost wages and reduced productivity exceeding $100 billion in the U.S. and $12 billion in the U.K. annually.[3,4] The most common diagnosis for patients experiencing LBP is intervertebral disc degeneration (IVDD),[5] which presents with several clinical characteristics. X-ray and magnetic resonance (MR) imaging often demonstrate significant reductions in overall IVD height, NP desiccation, intra-vertebral herniations (i.e. Schmorl nodes), changes in the vertebral body end-plates (i.e. Modic changes), and osteophyte formation.[6] These changes are associated with IVDD, an aberrant, cell-mediated process originating in the NP due in part to genetic predispositions, mechanical overload, and limited nutrient supply.[7] Once initiated, IVDD is marked by elevated concentrations of pro-inflammatory mediators that promote the production of matrix-degrading enzymes by resident cells.[8,9] In turn, matrix turnover will favor catabolism over anabolism, eventuating the desiccation and breakdown of the NP tissue; specifically, its proteoglycan constituents. Degraded NP matrix promotes inflammation, as evidenced by increased concentrations of pro-inflammatory cytokines including interlukin-1 beta (IL-1β) and tumor necrosis factor-alpha (TNF-α).[10] This positive-feedback cycle coupling inflammation and matrix degradation may amplify small perturbations in IVD physiology, ultimately leading to altered IVD microarchitecture and reduced matrix mechanical properties.[11] This begets altered spinal kinematics,[12] and non-physiologic loading of the AF which can result in lamellar disorganization and damage leading to herniation.[13]

The symptoms of IVDD are currently addressed by surgical discectomy to remove IVD tissue, joint fusion, or total disc replacement. These strategies suffer from significant limitations and may lead to degeneration of adjacent levels. To address these issues and restore IVD microarchitecture and function, regenerative medicine approaches are being investigated, including biologic administration and tissue engineering using novel biomaterial scaffolds and/or mesenchymal stem cells. Using such approaches, investigators have demonstrated the ability to promote IVD tissue regeneration *in vitro*,[14–16] as well as the attenuation of degeneration in small animal models.[17] However, clinical translation of these strategies requires evaluating their efficacy in large animal models that recapitulate the salient biochemical, mechanical, and clinical features of mild to moderate IVDD prior to human clinical trials.

Selecting the appropriate animal model to investigate IVDD and evaluate therapeutics has been controversial.[17,18] Reitmaier *et. al* reviewed large animal models used in IVD research and identified several shortcomings within the field, including minimal (e.g. based on available in vitro data) or no justification of animal model selection.[17] Additionally, the authors suggested that logistical factors, such as behavioral differences, animal cost, availability, or researcher preference, may influence model selection in the absence of empirical differences. The authors concluded that current limitations of data concerning large animal models minimized the impact of currently published literature, warranting further comprehensive characterization of *in vivo* models to determine their suitability for IVD research. Of those reviewed, caprine (goat) and ovine (sheep) models were found to be the most common quadrupeds used to study IVD pathology and repair.

In goats, Hoogendoorn and colleagues extensively characterized the ability of intradiscal injection of chondroitinase-ABC (C-ABC) to initiate and consistently produce progressive, degenerative changes in lumbar IVDs as evidenced by clinical imaging, biochemical, and histological analyses.[19–21] These results were re-affirmed in a similar study by Gullbrand *et. al* who administered 1U C-ABC which resulted in moderate IVDD in goats by 12 weeks.[22] In contrast, equally comprehensive studies have not been performed in sheep, despite significant similarities to human lumbar IVDs with respect to geometry,[23] range of motion,[18] matrix composition,[24] age-related changes,[25,26] notochordal cell absence,[27] and intradiscal pressures.[28] To date, only two studies have utilized intradiscal C-ABC injection to induce IVDD in this sheep.[29,30] Saski et. al performed intradiscal injections of 1, 5, or 50 U C-ABC per IVD, which resulted in significant reductions in intravital lumbar intradiscal pressures and heights over 4 weeks across all concentrations.[29] Ghosh et. al utilized 1U C-ABC to induce degeneration over a period of 12-weeks prior to administering progenitor cells to evaluate their efficacy to regenerate the IVD.[30] However, the study was not specifically designed to characterize the degenerative model and only included imaging and histological outcome analyses. Thus herein, a pilot study was undertaken to further characterize the biochemical, mechanical, and histological changes observed in sheep lumbar IVDs following intradiscal injection of 1U C-ABC and to compare the resulting degeneration to that observed in human pathology.

## Materials & Methods

The study was approved by the Institutional Animal Care and Use Committee of Colorado State University (IACUC Protocol: 16-6891A). Fifteen IVDs from five lumbar levels of three skeletally mature female sheep (*Ovis Aries*, Rambouillet ewes; 65-71 kg, 3 years-of-age, K&S Livestock, Fort Collins, CO) were utilized in this study. Animals were group housed in an indoor/outdoor pen with access to a three-sided shelter and evaluated daily by a veterinarian for signs of pain, behavior changes, or gait abnormalities for the duration of the study.

### Surgical procedure and intradiscal administration of C-ABC

Peri-operatively, a transdermal fentanyl patch (150 mcg/hr/sheep) was applied to each animal for five days starting one day prior to surgery. Twenty-four hours prior to surgery, five doses of procaine penicillin G (3 million units) and phenylbutazone (1 gram) was administered to each sheep. The animals were induced with ketamine (2mg/kg IV) and midazolam (0.2mg/kg IV) then intubated and maintained on 1.5-3% isoflurane in 100% oxygen throughout the surgical procedure. Using standard surgical preparation and aseptic technique, the lumbar IVDs were located via fluoroscopy. IVDD was induced via percutaneous intradiscal injection of 1U of C-ABC (Amsbio, Cambridge, MA) in 200 μL of vehicle (sterile 0.1% bovine serum albumin in 1× PBS - Fisher Scientific, Hampton, NH) into the L_1/2_, L_2/3_, L_3/4_ IVDs (n=9 IVDs) (Fig 1A). To confirm accurate needle position prior to injection, needle visualization was achieved using lateral and anterior-posterior fluoroscopy (Fig 1B). The L_4/5_ (n=3) and L_5/6_ (n=3) IVDs served as vehicle and uninjured controls, respectively. Post-operatively, sheep were monitored until ambulatory, and then returned to standard housing conditions. Following the progression of IVDD, driven by C-ABC injections, animals were euthanized by intravenous barbiturate overdose (pentobarbitone sodium, 88mg/kg) in compliance with the 2013 American Veterinary Medical Association guidelines. Immediately following euthanasia, AF and NP tissues from selected IVDs were immediately excised and frozen at −80°C for biochemical analysis. Lumbar spines were then harvested *en bloc* and shipped overnight on wet ice to Clemson University for further analysis.

**Fig 1.**
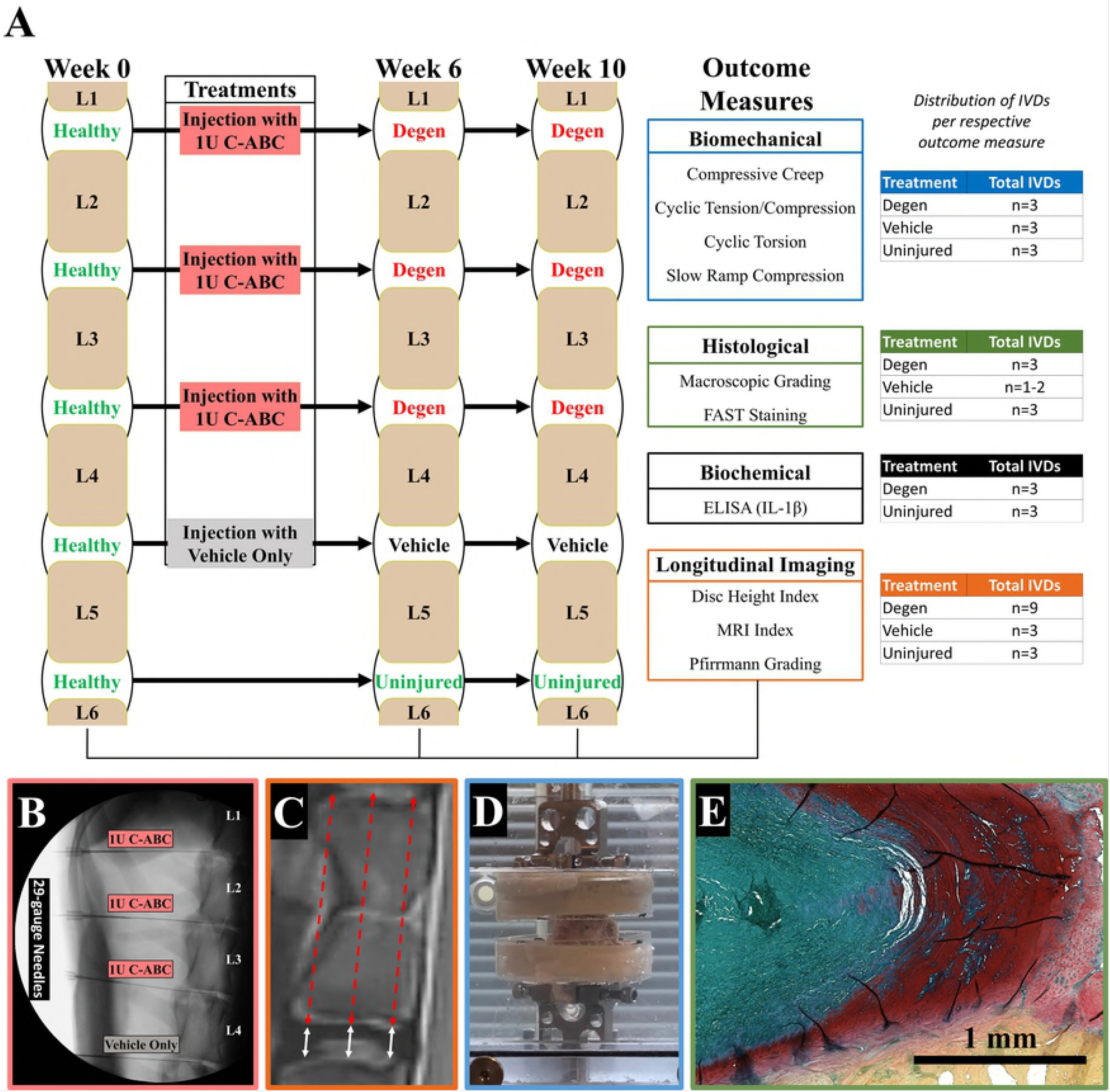
Animal study design overview and surgical approach. **A)** Animal study progressed over 10 weeks beginning with MR imaging and IVD degeneration (Degen) induction at week 0. At 6- and 10-weeks, IVDs were evaluated via MR imaging, kinematic testing, histological, and biochemical analysis. **B)** Representative lateral fluoroscopy image used to guide injections of 1U C-ABC in via 29-gauge needle. **C)** Representative T1-weighted MR image of an ovine vertebral body and adjacent IVD demonstrating IVD height (white solid arrows) and vertebral body height (red dotted arrows) measurements used for calculating DHI. **D)** Representative image of potted FSU undergoing kinematic creep loading in a saline bath. **E)** Representative histological image of an uninjured IVD stained with FAST: green /blue = NP, red = AF, and yellow = vertebral bone.

### Magnetic resonance imaging of IVDs

Sagittal MR imaging was performed immediately prior to intradiscal injection of C-ABC (week 0 – baseline) and tracked longitudinally at 6- and 10-weeks post-injection to monitor IVDD of degenerate (n=9), vehicle (n=3), and uninjured (n=3) IVDs. MR image scans were obtained using a 1.5 Tesla clinical imager (GE Signa). T2-weighted, T1-weighted, and Short-T1 Inversion Recovery (STIR) MR imaging sequences were performed on the explanted lumbar spines. Sagittal images were constructed using a T2-weighted fast spin echo sequence using a spine array coil (time to repetition: 2782 ms; time to echo: 101 ms; voxel size: 0.78 mm × 0.78 mm × 3.0 mm, with a 0 gap), a T1-weighted fast spin echo sequence using a spine array coil (time to repetition: 616 ms; time to echo: 18 ms; voxel size: 0.78 mm × 0.78 mm × 3.0 mm, with a 0 gap), and a STIR sequence using a spine array coil (time to repetition: 3500 ms; time to echo: 40 ms; voxel size: 1.56 mm × 1.56 mm × 3.0 mm, with a 0 gap and inversion time of 150).

Semi-quantitative MR image analysis was performed as described by Hoogendoorn et al.[21] MR imaging index was calculated from T2-weighted images as the product of the crosssectional area and mean signal intensity of the encircled NP using IMPAX 6.6.1.4024 (AGFA HealthCare N.V., Mortsel, Belgium) software. Consistent mid-sagittal IVD imaging was confirmed by ensuring the full cross-sectional of the spinal cord was in view. MR imaging index is expressed as a percentage of week 0 (pre-C-ABC injection) values for normalization. It should be noted that normalization of signal intensity was not performed by comparing to the spinal cord, as has been described by others, because it is a mobile structure with too much Gibbs and pulsation artifact. Thus, normalization to normal (week 0) IVDs was deemed most appropriate. Two researchers (R.B. and J.W.) quantified the MR imaging index for each IVD. A board-certified neuroradiologist (L.M.) verified the quantitation and performed qualitative analysis of MR images using the classification scale described by Pfirrmann et al.[31]

IVD height index (DHI) was measured from mid-sagittal, T1-weighted MR images in accordance with Hoogendoorn et al. with minor modifications.[21] IVD and vertebral body heights were each quantified using three height measurements at ventral-dorsal quartiles using ImageJ (NIH, Bethesda, MD) (Fig 1C). DHI was calculated as each IVD’s mean disc height divided by the mean adjacent vertebrae height, with changes expressed as a percent of week 0 DHI. Measurements were performed by two researchers (R.B. and J.W.) to evaluate interobserver reliability and blindly confirmed by board-certified neuroradiologist (L.M.).

### Enzyme Linked Immunosorbent Assay for Interlukin-1β

Ten weeks post-injection, AF and NP tissues from degenerate and uninjured IVDs (n=3/group) were assessed for the pro-inflammatory cytokine IL-1β. Tissues were cryo-pulverized and homogenized on ice in lysis buffer (500uL/10mg of tissue; 100mM TRIS buffer (pH=7.4), 150mM NaCl, 1mM EDTA, 1% Triton X-100, 0.5% Deoxycholic Acid, and 1× protease inhibitor cocktail (Sigma-Aldrich, St. Louis, MO)). Homogenates were then agitated for 60 min at 4°C, and the supernatant was isolated via centrifugation (5 minutes at 10,000×g). Supernatants underwent Enzyme-linked immunosorbent assay (ELISA) for ovine IL-1β (AB Clonal Science, Inc., Woburn, MA) according to the manufacturer’s instructions, the results of which were normalized to the total protein quantified via Bicinchoninic Acid (BCA) assay (ThermoFisher Scientific, Waltham, MA).

### Functional Spinal Unit Axial and Torsional Kinematics

Ten weeks post-injection, functional spinal units (FSU’s: vertebrae-IVD-vertebrae) of degenerate, vehicle, and uninjured (n=3/group) IVDs underwent biomechanical kinematic testing according to methods previously described by our group, with modification.[32] Briefly, FSU’s (with intact posterior elements) were potted in urethane resin (Goldenwest Manufacturing, Grass Valley, CA) (Fig 1D) first underwent creep loading on a Bose ElectroForce (model: 3220, TA Instruments, New Castle, DE) equipped with a 100-lb. load cell and a test chamber filled with 1xPBS/protease inhibitor at 25°C. Samples were loaded to −0.125 MPa and then underwent a 1-hr. creep period at −0.50 MPa. Samples were then transferred to a servohydraulic test frame (model: 8874, Instron, Norwood, MA) fitted with a 20kN load cell, and subjected to 35 cycles of axial compression (−0.50 MPa) and tension (0.25 MPa) at 0.1 Hz. Compression was then maintained at −0.50 MPa as samples underwent 35 torsion cycles to ±3°. Finally, samples underwent a slow-rate compressive ramp (1 N/s) from −0.125 MPa to −0.50 MPa. A non-linear constitutive model was fit to the creep data using GraphPad Prism 7 software (La Jolla, CA) to yield elastic (Ψ) and viscous (*η*) damping coefficients for the short-term (*η_1_* and Ψ_1_) and longterm (*η_2_* and Ψ_2_), as described previously.[33] Tensile and compressive stiffness was determined using a linear fit of the loading force-displacement curve from 60-100% of the 35^th^ cycle. Torsional stiffness was calculated from a linear fit of the loading torque-rotation curve of the 35th cycle. Torque range and axial range of motion (RoM) was calculated as the peak-to-peak torque and displacement, respectively. The constant-rate slow-ramp compression stiffness was determined using a linear fit of the slow-ramp load-displacement response.

### Macroscopic evaluations and histology of IVDs

Following mechanical testing, samples were excised from their pots and fixed for 7 days in 10% neutral-buffered formalin, followed by decalcification in 12% formic acid. Samples were cut into 3-mm sagittal slices and images were captured for macroscopic evaluation by 3 blinded observers (R.B., J.W., and J.M.) using the Thompson grading scale (n=3/degen and uninjured IVDs; n=2/vehicle IVDs).[34] These mid-sagittal slices were paraffin-embedded and 7μm sections were obtained; however, two of the three vehicle controls were lost during tissue processing. Staining was performed in accordance with methods form Leung et al.[35] Briefly, sections were rehydrated via ethanol gradations and stained with 3% Alcian blue (pH 1.0) for 8 min, 0.1% safranin-O for 6 min, 0.25% tartrazine for 10 sec, 0.001% fast green for 4 min. Micrographs were acquired using an Axio Vert.A1 microscope with AxioVision SE64 Rel. 4.9.1 software (Zeiss, San Diego, CA), and composite images were stitched together using FIJI analysis software.[36] Composite IVD micrographs (n = 3/group for degenerate and uninjured; n = 1 for vehicle) were scored by two blinded observers (R.B., J.M.) and an unblinded observer (J.W.) using the IVD degeneration scale described by Walter *et al*.[37] Endplate Integrity, AF Morphology, AF/NP Demarcation, NP Matrix Homogeneity, and NP Matrix Stain Intensity were each assigned scores from 0 to 2, and these were summed to produce a semi-quantitative aggregate score.

### Statistics

Results are represented as mean ± standard error of the mean (SEM) and significance was defined as (p≤0.05). Statistical analysis was performed using Prism 7 software. Comparisons were performed using a one-way ANOVA with Holm-Sidak method for multiple comparisons (MR imaging Index, DHI, ELISA) or Dunnett’s post-hoc (FSU kinematics and Thompson grading) analysis. Pfirrmann scoring was evaluated via a Kruskal-Wallis test with Dunn’s multiple comparisons. Assessment of inter-observer reliability of macroscopic Thompson grading was evaluated via Fleiss’ kappa statistic. Inter-observer reliability of histological scores was evaluated using IBM Statistical Package for Social Sciences (SPSS 24.0; IBM, Armonk, NY). The Intra-class Correlation Coefficient (ICC) was calculated using a two-way random model for absolute agreement as described by McGraw and Wong.[38] An ICC of 0.4-0.75 indicates good agreement, while >0.75 is considered excellent.[39] Mean histological scores were compared between degenerate and uninjured groups via one-tailed Mann Whitney test.

## Results

All animals tolerated the intradiscal C-ABC injection procedure, recovered from anesthesia, and began weight bearing and eating within one hour post-operatively. At 6 weeks, one animal had difficulty rising from a recumbent position with severe hind end weakness and moderate pain response upon palpation of the lumbar region. MR imaging of the animal’s lumbar spine revealed signs of severe IVDD and a large IVD protrusion / herniation into the spinal canal causing cord compression at the L_1/2_ level. The animal was euthanized, cultures and histopathology were obtained from the herniated IVD and the spine was harvested *en bloc*. No infection was found in the L_1/2_ IVD. Histopathology revealed reactive fibroplasia with mixed inflammation of lymphocytes, plasma cells, and macrophages. The remaining two degenerate IVDs (L_2/3_ and L_3/4_) were excluded from further analysis. The vehicle and uninjured IVDs (L_4/5_ and L5/6, respectively) were stored at −80°C, and later used for biochemical, biomechanical, and histological analysis. The remaining two animals recovered as expected without any signs of complication throughout the 10-week post-injection period.

### Intradiscal C-ABC injection results in significantly reduced MR imaging index and increased Pfirrmann grades of degeneration in IVDs

T2-weighted MR images illustrated a progressive darkening of the NP-region and adjacent vertebral endplates in degenerate IVDs (Fig 2A). Conversely, uninjured IVDs and those injected with vehicle demonstrated no such changes (Fig 2A). Of note, darkening in adjacent vertebral bodies adjacent to degenerate sheep IVDs were also observed on MR images indicative of endplate abnormalities (Fig 2A). Normalized MR imaging index of degenerate IVDs was significantly lower at week 6 (65.63±4.01%; p=0.024) and week 10 (57.10%; p=0.023) values. Moreover, these values were significantly lower compared to respective uninjured controls at week 6 (p=0.049) and continued to progress at week 10 (Fig 2B). No significant changes in MR imaging index were observed between uninjured and vehicle control IVDs. Degenerate IVDs also demonstrated a significant increase in Pfirrmann grade, indicating worsening degeneration at week 6 (2.11±0.261; p=0.014) and week 10 (2.833±0.401; p<0.001) compared to respective week 0 values (Fig 2C). Uninjured and vehicle control IVDs both had average Pfirrmann scores of 1±0 at all time-points investigated.

**Fig 2.**
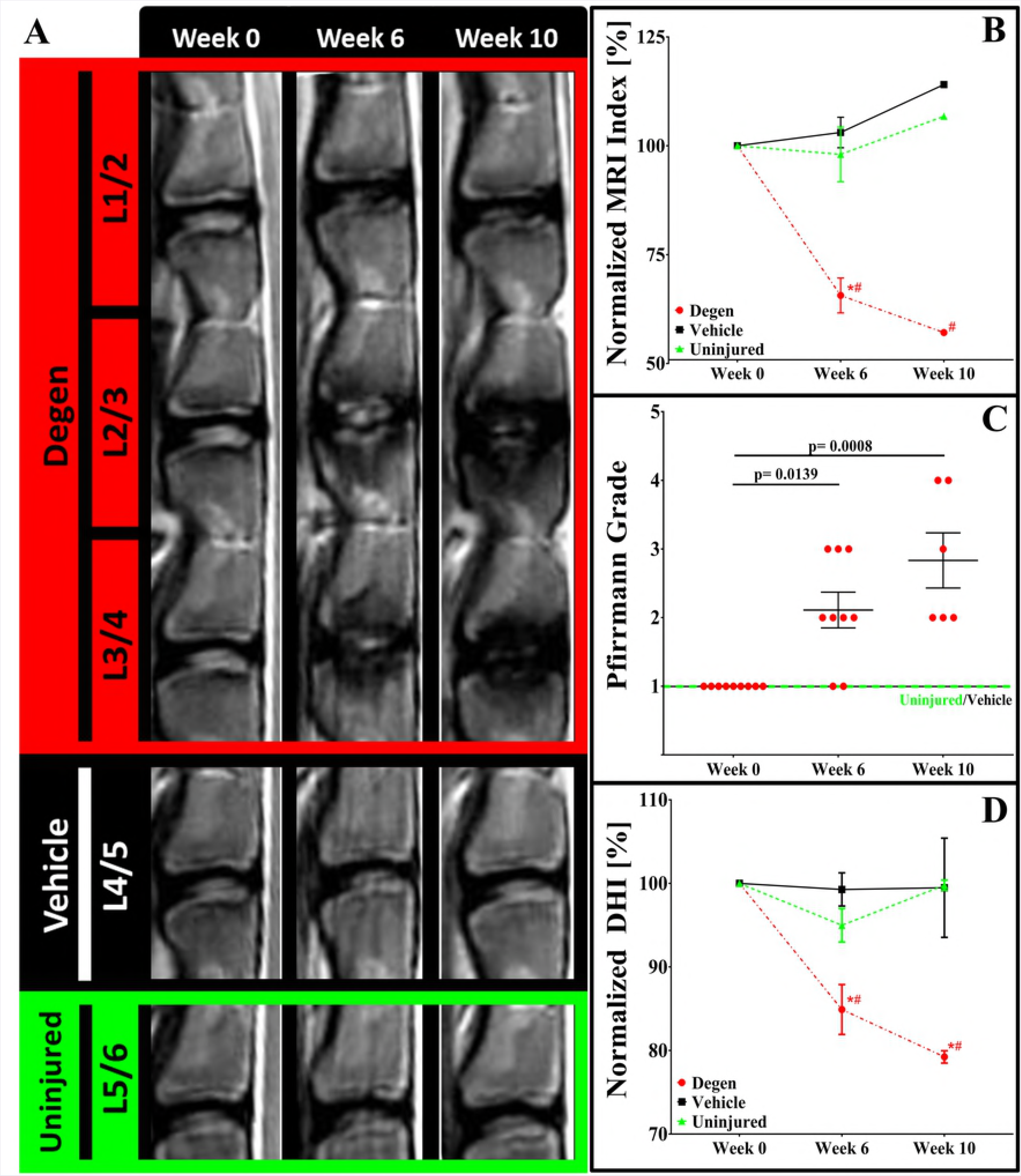
Longitudinal image tracking of sheep lumbar IVDs. A) Representative T2-weighted MR longitudinal images of a single sheep over the duration of the 10-week study. Graphs depicting quantitative MR image analysis for B) normalized MR image index, C) Pfirrmann grading, and D) normalized disc height index. (*) indicates a significant difference (p<0.05) between groups within the same time-point. (#) indicates a significant difference (p<0.05) within the study group compared across time-points. Solid lines connecting groups indicate a statistical difference (p<0.05).

### Intradiscal C-ABC injection results in significant reductions in IVD height

Normalized DHI values of degenerate IVDs significantly decreased over time, reaching 84.91±2.98% (p=0.007) at week 6 and 79.23±0.74% (p<0.001) at week 10 (Fig 2D). These values were also significantly lower compared to uninjured IVD values at week 6 (94.99±2.00%; p=0.048) and week 10 (99.78±0.61%; p= 0.002), respectively. No significant changes in DHI were observed between uninjured and vehicle control IVDs.

#### Intradiscal C-ABC injection results in significantly increased concentrations of IL-1β

ELISA analysis of degenerate IVDs demonstrated a significant (p=0.002) increase in pro-inflammatory IL-1β concentrations compared to uninjured IVDs (483.2±79.0 [pg/mL]/mg vs. 39.7±34.1 [pg/mL]/mg, respectively) (Fig 3). IL-1β was found to be increased in both the NP- and AF-regions of degenerate IVDs compared to uninjured IVDs; however, this was significant (p=0.019) only in the NP (Fig 3).

**Fig 3.**
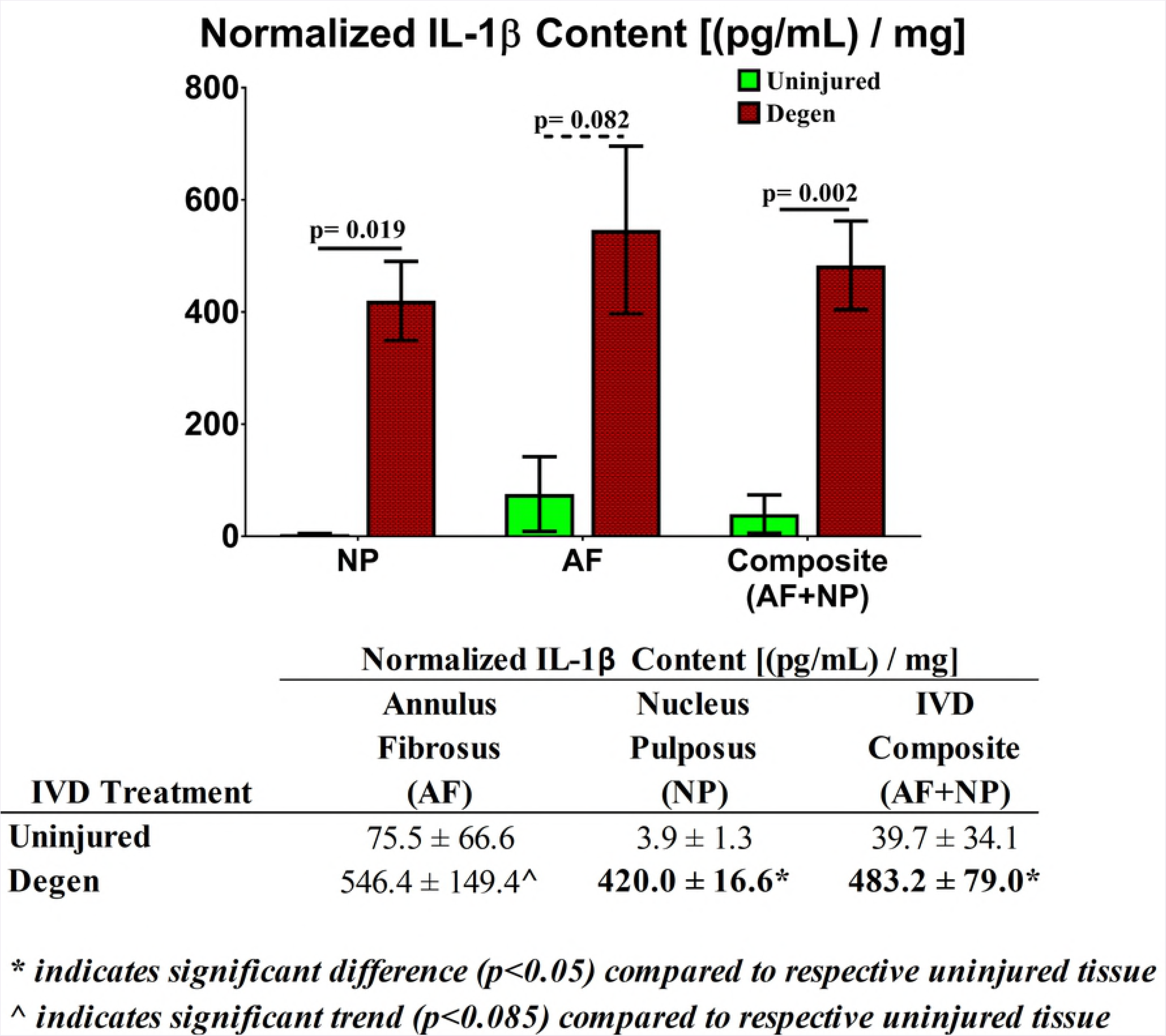
IL-1β quantification of sheep IVDs. Graphical and tabular results of normalized IL-1β concentrations from the NP and AF regions of degenerate and uninjured sheep IVDs. Solid lines connecting groups indicate a statistical difference (p<0.05). Dotted lines indicate a trend towards significance (p<0.085).

### Intradiscal C-ABC injection results in significant alterations in functional spinal unit kinematics

Kinematic loading (Fig 4A) of FSUs containing degenerate IVDs demonstrated a reduced step displacement (0.11±0.02 mm) and significantly (p=0.022) increased creep displacement (0.50±0.05 mm) compared to FSUs with uninjured IVDs (0.16±0.016 and 0.30±0.02 mm, respectively) (Fig 4C). Additionally, long-term elastic (Ψ_2_: 482.17 ± 69.89 N/mm) and viscous (η_2_: 92.1 ± 2.78 × 10^4^Ns/mm) damping coefficients of FSUs with degenerate IVDs were significantly (p=0.0074 and p=0.0016, respectively) lower compared to FSUs with uninjured IVDs (Ψ_2_: 862.26 ± 29.28 N/mm and η_2_: 229.52 ± 17.86 ×10^4^Ns/mm, respectively) (Figs 4D and 4E).

**Fig 4.**
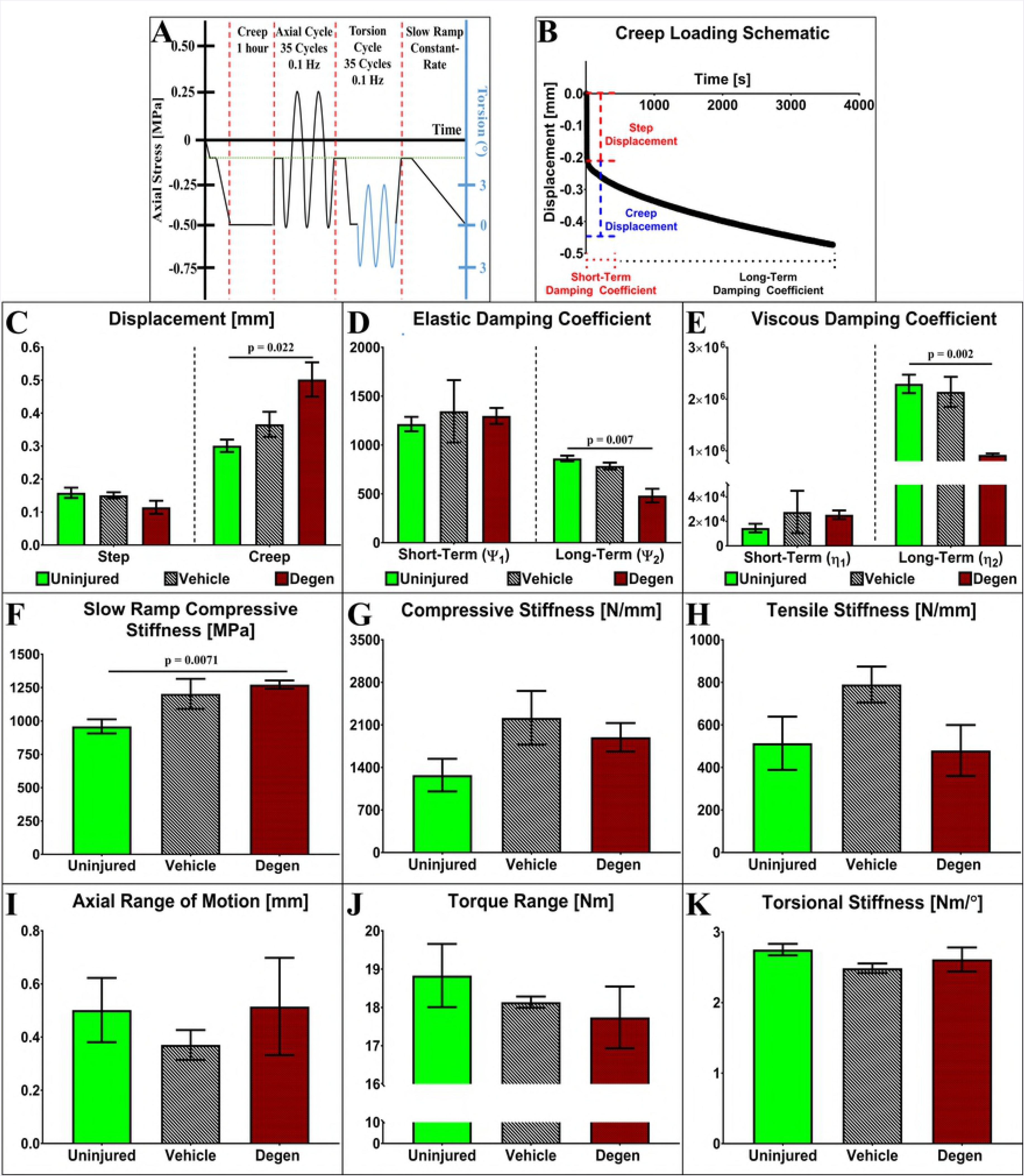
Kinematic testing of IVD FSUs. **A)** Loading scheme for FSU testing depicting creep, axial cyclic tension-compression, axial torsion, and slow constant-rate ramp testing. **B**) Representative graph depicting the creep response of a sheep IVD and associated creep parameter measures. Graphs depicting **C)** step and creep displacement, **D)** elastic damping and **E)** viscous damping coefficients, **F)** slow ramp compressive stiffness, G) compressive stiffness, **H)** tensile stiffness, **I)** axial RoM, **J)** torque range, and **K)** torsional stiffness values of uninjured, vehicle and degenerate (degen) sheep IVDs, respectively. Solid lines connecting groups indicates significant difference (p≤0.05) compared to uninjured controls.

Slow-ramp compressive loading demonstrated a significant increase (p=0.007) in compressive stiffness of FSUs containing degenerate IVDs compared to those with uninured IVDs (1272.00±31.76 MPa and 959.4±52.87 MPa, respectively) (Fig 4F). A similar trend was observed for cyclic compressive stiffness of FSUs containing degenerate IVDs (Fig 4G). No significant changes were observed in the axial tensile stiffness or range of motion (Figs 4H and 4I). Torsional rotation of FSUs containing degenerate IVDs demonstrated a decrease in torque range (17.74±0.81 Nm vs. 18.83±0.82 Nm, respectively) and torsional stiffness (2.61±0.17 Nm/° vs. 2.75±0.08 Nm/°, respectively) compared to uninjured IVDs (Figs 4J and 4K). No significant changes were observed within the kinematic parameters between FSUs containing uninjured and vehicle control IVDs.

### Intradiscal C-ABC injection induced significant changes in IVD morphology and microarchitecture

Macroscopic evaluation of IVDs using the Thompson scale (Fig 5) showed substantial agreement for inter-observer variability (κ grade: 0.75) and showed that the average Thompson grades for degenerate IVDs (3.89±0.20), were significantly greater compared to uninjured (1.00±0.00; p<0.001) and vehicle controls (1.00±0.00; p<0.001), respectively.

**Fig 5.**
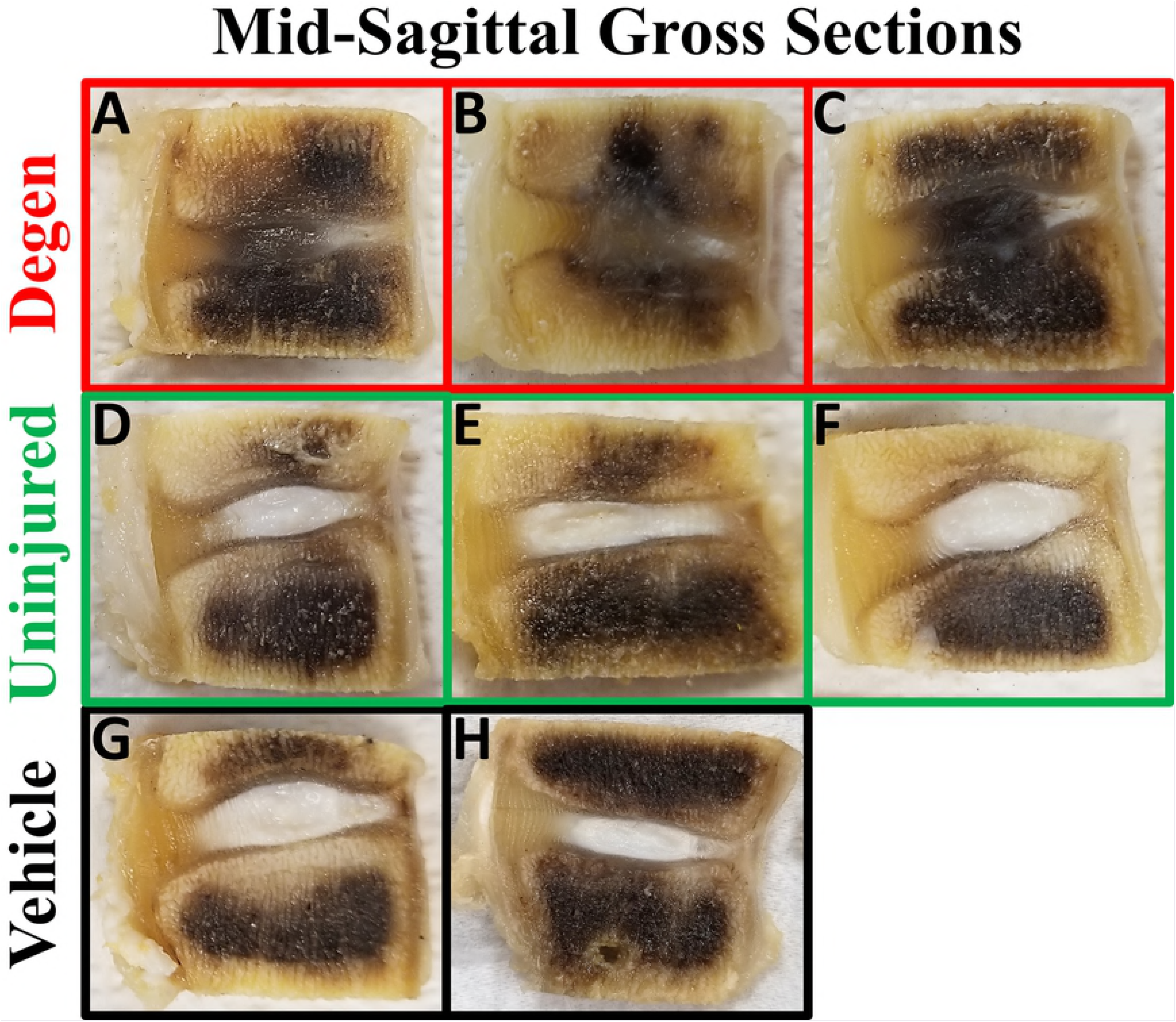
Mid-sagittal gross sections of ovine IVDs. Representative IVDs utilized for Thompson grading depicting the macroscopic changes between **A-C)** 1U C-ABC injected IVDs (degen), **D-F)** uninjured IVDs, and **G-H)** vehicle IVDs.

Semi-quantitative histological scoring showed excellent agreement among observers (ICC: 0.975) and significant differences for uninjured and degenerate scores. (Fig 6) Qualitatively, degenerate IVDs revealed significant alterations and disruption of tissue architecture compared to control IVDs (Figs 6A-F). Prominent intravertebral herniations were observed in all evaluated degenerate IVDs (Figs 6D-F), and these consistently exhibited several abnormalities compared to control IVDs: 1) failure and displacement of the CEP into the subchondral bone, 2) extrusion of marked amounts of NP into the vertebrae, 3) apparent remodeling around the herniated tissue, and 4) thickening of the vertebral bone. The AF of degenerate IVDs also showed a notable change in concavity, in contrast to the control IVDs that showed normal, convex AF orientation (Fig 6). Furthermore, gross morphological changes in degenerate IVDs seemed to have pronounced effects on AF structure. Specifically, loss of IVD height seemed to compress AF lamellae while CEP displacement resulted in reduced convexity. Higher magnifications revealed changes in distinct regions of the IVD (Fig 7). The ability of the CEP and NP to retain stain was severely diminished by C-ABC injection. Additionally, thinning and permeabilization of the CEP was seen through microscopic NP expulsion and reduction in CEP thickness. The vehicle control showed no qualitative histological differences compared to the uninjured samples and received a comparable aggregate score (0.6±0.4 for uninjured vs 0 for vehicle).

**Fig 6.**
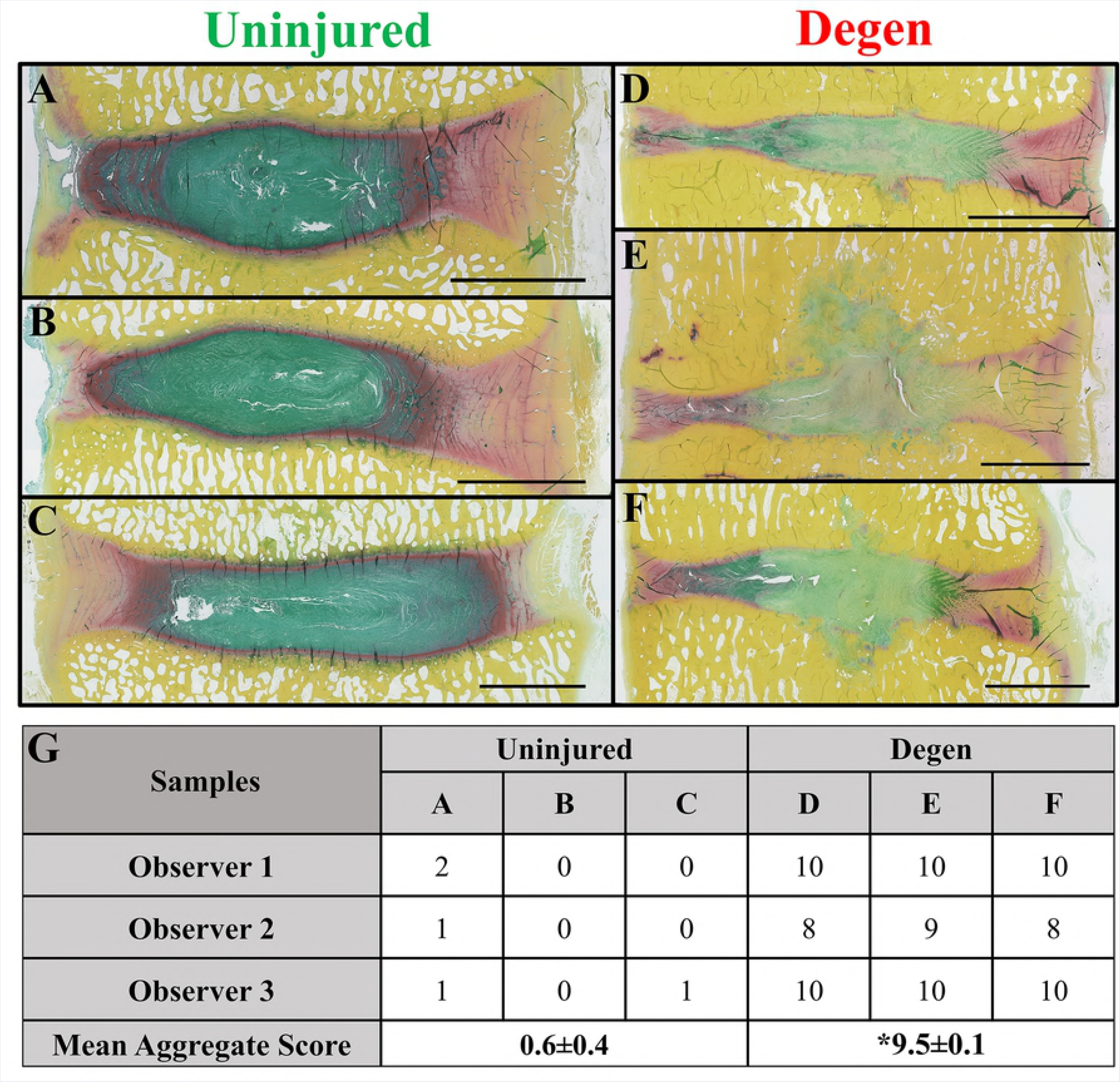
Semi-quantitative scoring of ovine lumbar IVDs. Macroscopic images of **(A-C)** uninjured and **(D-F)** degen IVDs were used for scoring. **G)** Aggregate scores for uninjured and degenerate IVDs were calculated as the average of each observer and sample score within the respective treatment. Significance is indicated with ‘*’ (p≤0.05). Scale bar = 5 mm. *FAST Staining:* Greenish-blue = glycosaminoglycan / NP, Red = CEP/AF, Yellow = vertebral bone, and Light Green = outer AF.

**Fig 7.**
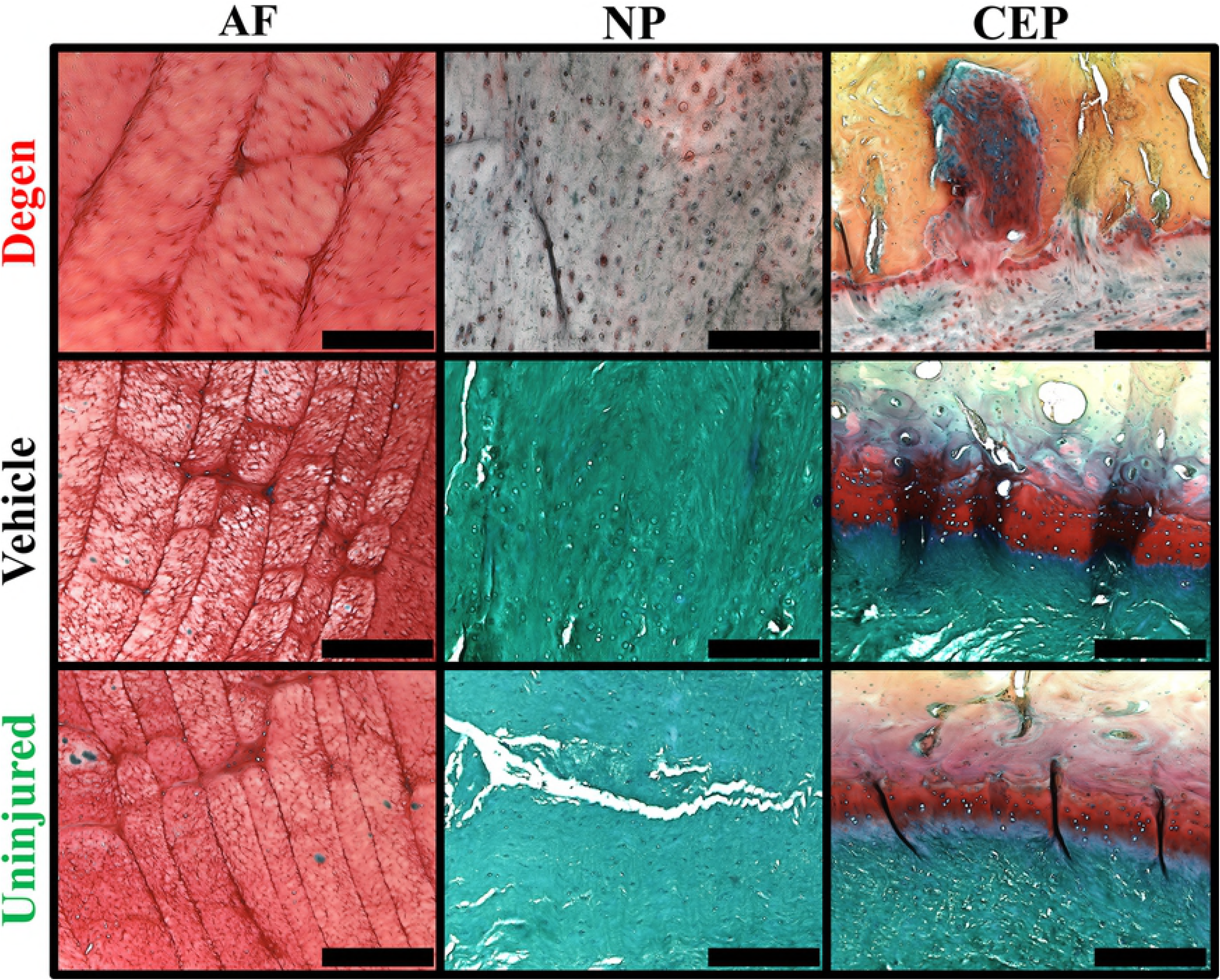
Qualitative histological comparison of anatomical regions of the IVD across study groups. **Left column**, representative histological images of AF lamellae from degenerate, uninjured and vehicle control IVDs depicting differences in lamellar thickness (red = AF lamellae). **Center column**, representative histological images of the NP region of degenerate, uninjured and vehicle control IVDs illustrating difference in glycosaminoglycan staining intensity (blue/green = glycosaminoglycan). **Right column**, representative histological images of the cartilaginous end-plates of degenerate, uninjured and vehicle control IVDs illustrating differences in integrity and thickness. Scale bar = 400 μm. *FAST Staining:* Greenish-blue = glycosaminoglycan / NP, Red = CEP/AF, Yellow = vertebral bone, and Light Green = outer AF.

## Discussion

The present pilot study aimed to build upon the initial work of Saski et. al and Ghosh et. al who demonstrated that intradiscal administration of C-ABC resulted in mild to moderate IVDD as evidenced by reductions in IVD intradiscal pressure, height, hydration, and changes in microarchitecture.[29,30] Two-hundred microliter intradiscal injections of 1U C-ABC were performed in the lumbar spine of three sheep and degenerative changes were tracked for up to 10 weeks in alignment with previous studies by others.[22,29,30] The primary findings of this pilot study herein suggest that intradiscal injection of 1U C-ABC in sheep resulted in moderate to severe IVD degeneration marked by progressive and significant alterations in IVD hydration, height, inflammation, kinematics, and IVD tissue morphology (including the formation of end-plate disruptions). The observed changes appeared to be potentially more severe as compared to previous reports;[20–22,29,30] however, these changes were still reminiscent of that observed in human lumbar IVDD.

In the present pilot study, clinical imaging of sheep C-ABC injected IVDs demonstrated significant reductions in hydration and height, which worsened over time. This was evidenced by progressive decreases in both T2-weighted MR image signal intensity and IVD height measurements. No evidence of spontaneous regeneration was observed during the study time-course. Such observations are likely due to a loss of aggregating proteoglycan, leading reduced NP osmotic potential, lowered water content, and the apparent increase in matrix permeability.[40–43] Together, these alterations diminish the ability of the tissue to effectively support compressive loading leading to a reduction in IVD height and overall mechanical dysfunction.[6,11,44] Similarly, imaging of degenerate human IVDs often display progressive darkening of the IVDs and loss of height with increasing grade of degeneration.

Trends in the Pfirrmann grading scores for degenerate IVDs reflected macroscopic deterioration, with some IVDs receiving scores of 3-4. This is indicative of demarcation between the NP and AF regions of the IVD in conjunction with reductions in MR image signal intensity and IVD height.[31] From a clinical perspective, IVDs having a Pfirrmann grade 3 or lower remain functional and would be targets for regenerative medicine approaches employing biomaterials, biologics, and cells. Notably, MR imaging corroborated the presence of subchondral end-plate irregularities adjacent to degenerate IVDs as evidenced by the darkening and obscuring of these structures. Likewise, patients with lumbar IVDD often exhibit similar findings which are referred to as Modic changes.[45] Such changes represent a spectrum of biological alterations occurring within vertebral bodies which can include fatty marrow changes, end plate sclerosis, trabecular fracture, as well as edema and inflammation.[45]

Biochemical analysis of degenerate sheep IVDs demonstrated an increased concentration of IL-1β compared to control samples, indicating elevated inflammation. These findings are likely related to the observed endplate changes discussed above. Degenerate human lumbar IVDs have been shown to contain increased concentrations of pro-inflammatory cytokines, including IL-1β, which are produced by endogenous IVD cells and infiltrating inflammatory cells.[10] The increased presence of this cytokine has been shown to result in the up-regulation of matrix degrading proteases by NP and AF cells, which further compromise the extracellular matrix (ECM) and mechanical function of the IVD.[46–48]

Axial and torsional kinematics of degenerate sheep IVDs were significantly altered compared to uninjured and vehicle control samples. Most notably, the creep displacement significantly increased in conjunction with significant reductions in the long-term elastic and viscous dampening time constants. Together, this indicates that the ECM of the IVD has been disrupted and is no longer able to as effectively resist compressive creep loading as compared to controls. This is likely an effect of degradation of the ECM in conjunction with an increase in its permeability leading to reduced pressurization. Interestingly, the compressive stiffness of the degenerate sheep IVDs was significantly greater than uninjured controls. This has been observed by others and could be due to a reactive process involving the posterior elements in response to inflammation and overloading, and thus are attempting to stabilize the degenerate FSU’s.[49] This could also explain why overall FSU range of motion was ultimately unaltered in the degenerate study group. Similar phenomena have been observed in human degenerate lumbar IVDs in which spinal instability (i.e. increase in neutral zone and overall range of motion) occurs in early- and mild degeneration,[50,51] however restoration or re-stabilization of axial spinal kinematics has been shown to occur as degeneration progresses.[49] However, further study of this in the sheep model is warranted.

Semi-quantitative and qualitative histological evaluations of degenerate sheep IVDs demonstrated significant alterations in the NP, AF, and endplate micro-architecture as compared to controls. This was demonstrated by decreases in proteoglycan staining intensity, alterations in lamellar architecture, and the formation of Schmorl nodes, respectively. Taken together, these histomorphological features are suggestive of moderate to severe IVDD. We hypothesize that C-ABC degraded NP proteoglycan and the cartilaginous endplates. In fact, cartilaginous endplate thickness was found to be irregular in both macroscopic (i.e. Thompson scoring images) and microscopic analysis. Loss of cartilaginous endplate integrity due to C-ABC treatment could account for the observed formation of Schmorl nodes and the resultant increase in inflammation. The cartilaginous endplate functions to prevent the NP from breaching the vertebral body which prohibits its contact with bone marrow and the immune system. This is critical as NP tissue has been shown to be immunogenic when it herniates extradiscally.[52] It is plausible that because sheep lumbar IVDs have been shown to have higher intradiscal pressures compared to human IVDs,[17,53] in combination with enzymatic disruption of the cartilaginous endplate, most likely resulted in NP tissue breaching into the vertebral body. Such breaches were not observed in vehicle control IVDs which suggests that enzymatic damage to the endplate as opposed to an increase in intradiscal pressure due to injection volume caused the observed Schmorl node formation. Additionally, the formation of Schmorl nodes could be due to alterations in the vertebral body bone as a result of inflammatory edema due to breach of the cartilaginous end-plate and exposure of the NP,[54] however this was not directly assessed in the present study. In the context of lumbar IVDD in humans, the presence of Schmorl nodes have been shown to be correlated with Modic changes and grade of degeneration in human IVDs.[54,55]

As with any study, limitations were noted. First, this pilot study used a small sample size (n=3 sheep) to evaluate the effects of intradiscal injection of C-ABC on IVDD. Despite this fact, resultant degeneration was severe enough to cause statistical differences in many of the outcomes measured. Moreover, the promising results of this pilot study provides the foundation for us to continue to investigate this ovine model using a larger sample size and additional outcome measures: including, changes in IVD GAG and collagen content, in addition to, quantifying several pro-inflammatory mediators and proteases. Second, while we used an established T2-weighted MR imaging modality to evaluate ECM degradation as a function of IVD hydration, it would be advantageous to include a T1rho relaxation parameter, which may correlate more strongly with changes in IVD GAG content and histology.[56] Third, IVDs receiving C-ABC injection and those serving as controls were not randomized to account for differences in spine level. However, data from each IVD was normalized to its respective week 0 (baseline) value to account for this.

## Conclusion

In conclusion, we have begun the in-depth characterization of a chemonucleolysis-induced model of IVDD in sheep. Intradiscal administration of 1U C-ABC per lumbar IVD resulted in many of the salient hallmarks observed in the human pathology. Although further study is warranted using a larger sample size, this model may represent a valuable tool in the future for evaluating strategies to attenuate IVDD and its resultant sequelae.

## Acknowledgements

The authors would like to acknowledge the veterinary staff at Colorado State University, Dr. Patrick Gerard for his assistance with statistical analysis, and the Greenville Health System for allowing us to use their IMPAX Radiology Information System.

## Author contributions

Study design (R.B., J.W., S.G., J.M., L.M.), surgical implementation (J.E.), sample processing (R.B., J.W., A.M.), data analysis (R.B., J.W., J.M., A.M., L.M.), manuscript preparation (R.B., J.W., J.M.). All authors have read and approved the final submitted manuscript.

